# Fluorescence signatures of SARS-CoV-2 spike S1 proteins and an human ACE-2: excitation-emission maps and fluorescence lifetimes

**DOI:** 10.1101/2021.05.20.444935

**Authors:** Jonas Grzesiak, Lea Fellner, Karin Grünewald, Christoph Kölbl, Arne Walter, Reinhold Horlacher, Frank Duschek

## Abstract

**Significance:** Fast and reliable detection of infectious SARS-CoV-2 virus loads is an important issue. Fluorescence spectroscopy is a sensitive tool to do so in clean environments. This presumes a comprehensive knowledge of fluorescence data.

**Aim:** This work aims at providing fully featured information on wavelength and time-dependent data of the fluorescence of the SARS-CoV-2 spike protein S1 subunit, its receptor binding domain (RBD) and the human angiotensin-converting enzyme 2 (hACE2), especially with respect to possible optical detection schemes.

**Approach:** Spectrally resolved excitation-emission maps of the involved proteins and measurements of fluorescence lifetimes were recorded for excitations from 220 to 295 nm. The fluorescence decay times were extracted by using a bi-exponential kinetic approach. The binding process in the SARS-CoV-2 RBD was likewise examined for spectroscopic changes.

**Results:** Distinct spectral features for each protein are pointed out in relevant spectra extracted from the excitation emission maps. We also identify minor spectroscopic changes under the binding process. The decay times in the bi-exponential model are found to be (2.0*±* 0.1) ns and (8.0 *±*1.0) ns.

**Conclusions:** Specific material data serve as important background information for the design of optical detection and testing methods for SARS-CoV-2 loaded media.

## 1 Introduction

The global spread of the SARS-CoV-2 virus, which started in early 2020 [1] and which was declared pandemic by the WHO in March 2020 [2], has promoted the intensive research on human coronaviruses [3], which are a global public health thread [4]. A major tool in monitoring and controlling the spread of the viruses is a fast, accurate and sensitive detection of the virus or the infection[5]. In the case of SARS-CoV-2, a significant number of research articles on sampling techniques, nano biosensor technologies and antigen testing have been published and are reviewed e.g. in Refs. [6], [7, 8] and [9], respectively. An overview on photonic approaches for detecting virus loads and infections by optical means is given in Ref. [10]. The authors summarize that spectroscopic techniques, such as FTIR, Raman and fluorescence, are still facing challenges from the huge diversity of viruses in the human organism. Among those methods, laser induced fluorescence (LIF) is a widely used tool to investigate proteins [11, 12, 13, 14]. Elastic light scattering and fluorescence have been used to observe virus particles and virus-like particles [15, 16, 17]. The LIF technology takes advantage of increased signal intensities and has been applied to detect and classify viruses with excitation in the near UV [18, 19]. As pointed out in Ref. [20], additional, orthogonal information on fluorescence characteristics can be retrieved not only from two-dimensional, spectral signatures (excitation-emission maps) but also from fluorescence decay times. Such distinct features are of high importance for a reliable sample classification[21].

In this work, high resolution fluorescence data of the proteins involved in the SARS-CoV-2 binding process, namely the S1 part of the spike protein of SARS-CoV-2, the receptor-binding domain thereof and human angiotensin-converting enzyme 2 (hACE2) are reported as well as results on their fluorescence decay times. Spectral data and fluorescence decay times of the investigated proteins are compared and discussed as candidates for classification features.

## 2 Experimental setup

The proteins used for this study are SARS-CoV-2 S1 RBD (S1 RBD domain with His-Tag, M=27.5 kDa), SARS-CoV-2 S1 (S1 domain with His-tag, 77.1 kDa) and hACE2 (ECD domain, tag free processes M=80 kDa). The proteins were produced by trenzyme GmbH using a transient production system in HEK293 suspension cells followed by purification either via the encoded His-Tag by IMAC (RBD and S1 protein) or by ion exchange chromatography (hACE2). After purification, the buffer of the purified proteins was exchanged to DPBS (Dulbecco’s phosphate buffered saline (DPBS), pH=7.1-7.5, Sigma-Aldrich) and the proteins were analyzed. The final concentrations of the solutions were 22.6*µ*g*/*ml in case of the hACE2, 7.8*µ*g*/*mlin the RBD case and 22.2*µ*g*/*ml for the full length S1 protein. This ensures comparable, molar concentrations of the proteins for the analysis.

The samples were investigated in UV-transparent cuvettes (fused silica glass, Hellma 117F), with a light travel path of 10 mm in the spectrometer.

A FS5 fluorescence spectrometer from Edinburgh Instruments was used to obtain full fluorescence signatures of the proteins. The spectrometer uses a 90^*°*^ setup and a variable wavelength excitation. Additionally it allows for time correlated single photon measurements of the samples. The fluorescence maps contain data for excitation wavelengths ranging from 220 nm – 300 nm and emission from 230 nm – 500 nm. Time dependent fluorescence signals have been recorded at an excitation at 260 nm, with 0.9 ns excitation pulse duration and the respective optimal emission wavelengths.

## 3 Results and Discussion

### 3.1 LIF spectral signatures of the hACE2 enzyme and the S1 protein

As an overview, the emission maps of the SARS-CoV-2 S1 spike protein, the receptor-binding domain (RBD) thereof and the hACE2 are shown in Fig. 1. The diagonal trace of peaks, at +3430cm^−1^ from the excitation wavelengths *λ*_*ex*_ are assigned to be the Raman signals of H_2_O. The narrow peaks in the lower right corner of the graphs are assigned to be the second order diffraction maximum. Each map consists of two major maxima: the first one located at an excitation wavelength *λ*_*ex*_ = 220 nm with maximum emission at *λ*_*em*_ =325 nm for S1 and hACE2 and at *λ*_*em*_ = 320 nm for the receptor-binding domain. The second major maximum appears at an excitation wavelength of *λ*_*ex*_ = 280 nm showing a maximum emission at *λ*_*em*_ =325 nm for all three proteins.

**Fig 1.**
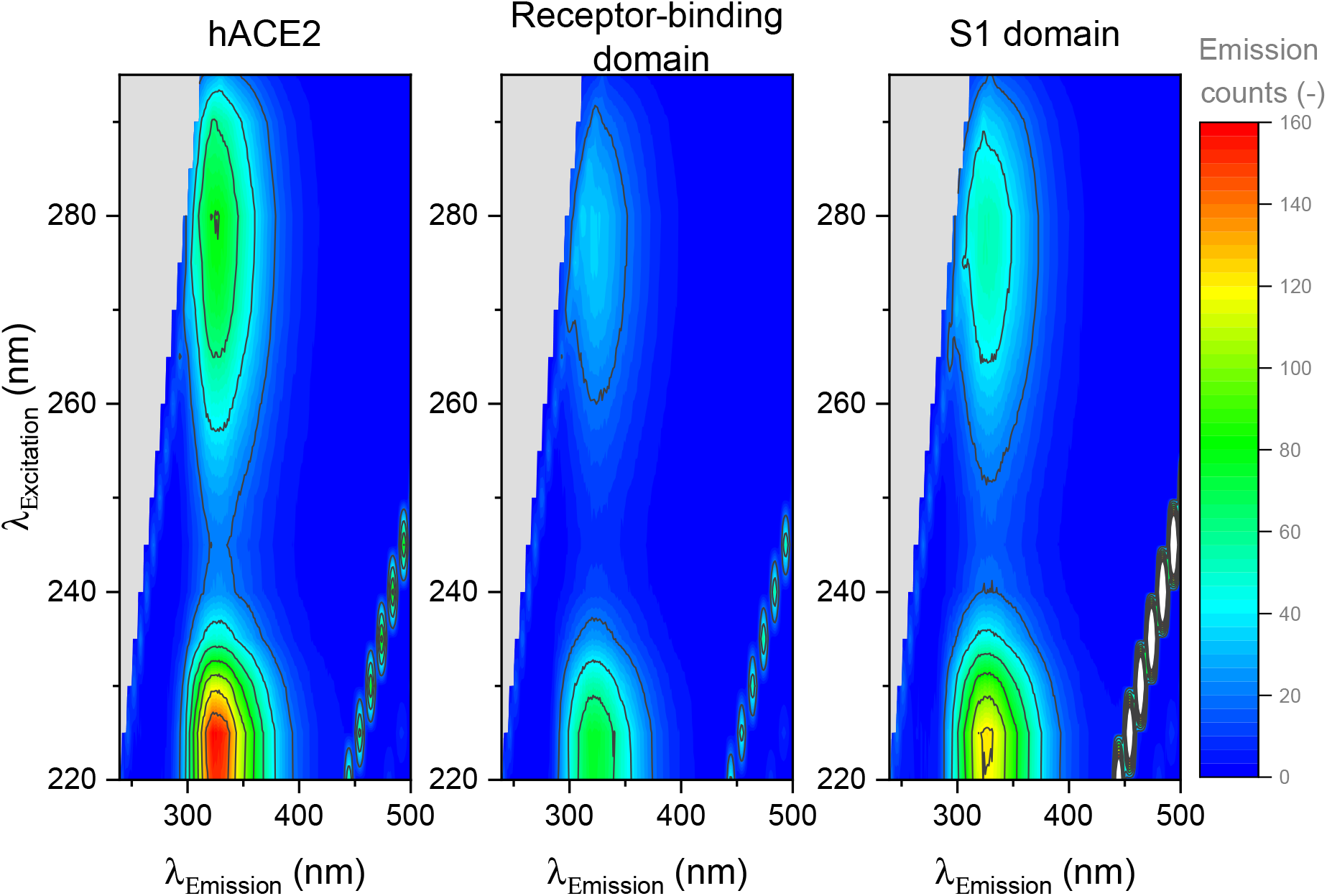
Excitation-emission fluorescence map for hACE2 (left), the receptor-binding domain (center) and SARS-CoV-2 S1 (right). Data are available on biorxiv.org as supplementary material.

Fig. 2 displays the normalized fluorescence response of the three samples in detail at excitation wavelengths *λ*_*ex*_ = 220 nm and *λ*_*ex*_ = 280 nm. For clearness the emission spectra obtained at *λ*_*ex*_ = 280 nm have been adjusted by the background signal from a pure PBS sample. The signals at *λ*_*ex*_ = 220 nm clearly show the Raman signal of the solvent as well as the second order diffraction maximum. The intensities of the respective signals are indicated by the Raman signal intensities and are more explicit in Fig. 3. The fluorescence peaks are located between 290 nm and 410 nm, which indicates that the excitation light mainly probes tryptophane and tyrosine sites inside the proteins. At both respective excitation wavelengths, the three proteins show significantly different fluorescence responses. This is manifested in differently shaped peaks, which in turn give full width half maximum values of the peaks from 55 nm for hACE2 and the receptor-binding domain to 60 nm for the S1 protein, both at an excitation wavelength of *λ*_*ex*_ = 280 nm. The same pattern can be seen for *λ*_*ex*_ = 220 nm, with slightly broader values for full width at half maximum (FWHM): 58 nm for hACE2 and RBD and 63 nm for the S1 protein. These slightly broader spectra may emerge from an excited hot (i.e. vibrationally excited) state with broad emission spectrum due to the lower excitation wavelength.

**Fig 2.**
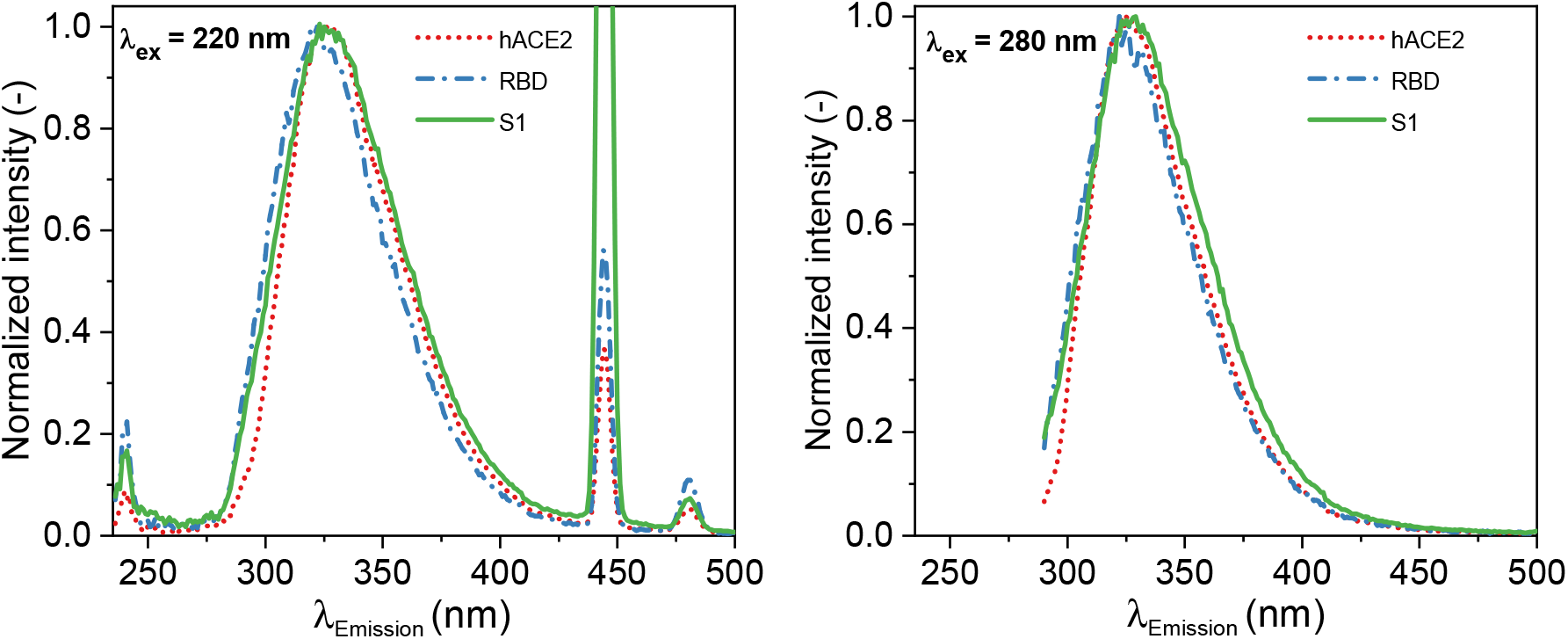
Detailed, normalized fluorescence signals of the three proteins at *λ*_*ex*_ = 220*nm* (left) and *λ*_*ex*_ = 280*nm* (right).

**Fig 3.**
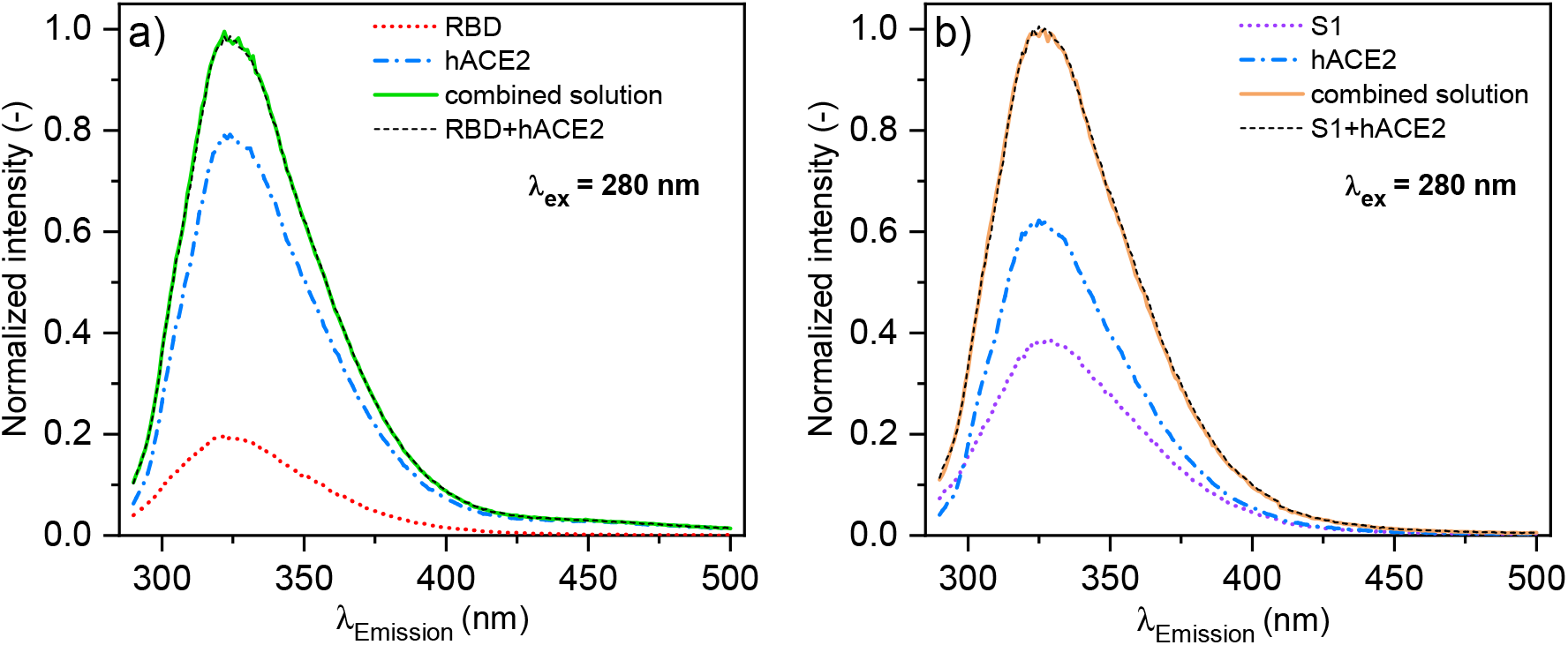
Fluorescence spectra of bound protein complexes and their single contributions: a) Receptor-binding domain of the S1 protein and b) the full S1 protein.

Besides the pure protein signals, also the bound complex of the virus with its receptor is of interest for a possible virus detection. To obtain a protein-enzyme complex solution we combined both the solution of the S1 full length protein and the RBD protein solution with the hACE2 solution: 1 ml of each solution with the respective concentrations noted above to ensure equimolarity of both substances in the combined solution. Straight after combination, the mixture was stirred to ensure a fully mixed solution. From the receptor binding affinity of about 1-40 nM [22] and the concentration of the S1 protein and the RBD protein analytes of about 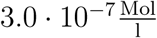 one obtains a half life time of the binding process in the range of *τ ≈* 3 − 35 s. Thus, the full binding of all hACE2 receptors to the virus proteins in our solution is ensured within 5 min, when the spectra were taken. Fig. 3 shows the fluorescence results of the bound complexes at an excitation wavelength of 280 nm. The signals have been adjusted by the background signal from a pure PBS sample and for simplicity the normalized signals are shown. For comparison the single contributions are shown too. We compared the signal of the bound complexes with the sum of the signals of the single contributions (black dashed lines in the spectra). The difference spectra between the summed signals and the bound complexes are shown in Fig. 4. Here, the significant change of about -1% of the signal at 312 nm is due to differences in the H_2_O Raman signal. At 350 – 400 nm however, a slight change in the slopes of the peak is found, indicating a possible conformation change under the binding process, especially in the S1 case.

**Fig 4.**
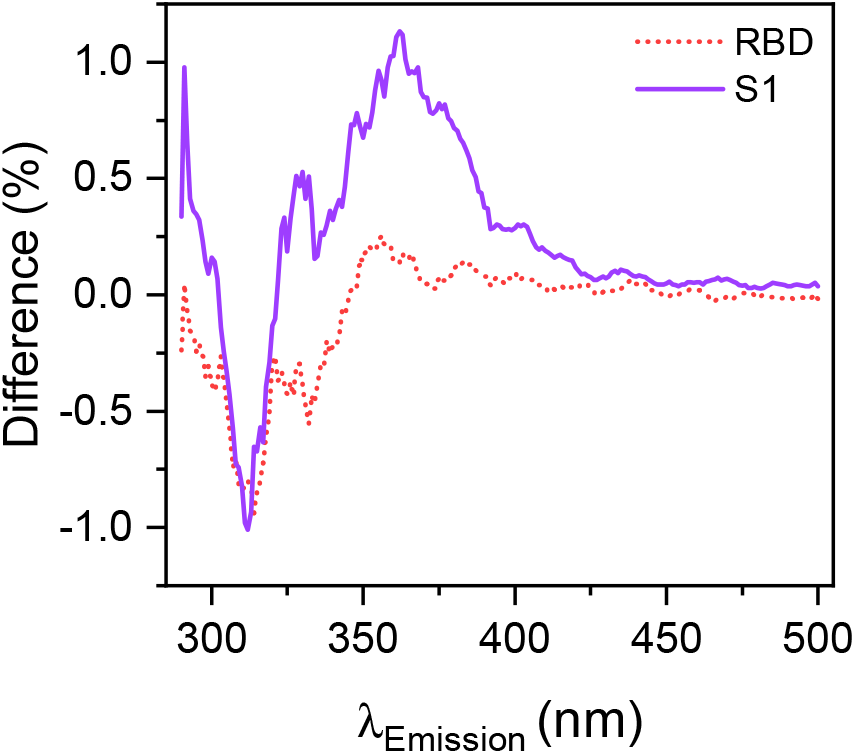
Differences between the added single contributions and the bound complex for the RBD and the S1 case.

### 3.2 Fluorescence decay times

Additionally, the fluorescence decay of the base solutions of the hACE2, the RBD protein and the S1 protein as well as of their bound complex solutions were measured at an excitation of 260 nm as a time correlated single photon counting signal, shown in Fig. 5. The decay signals were evaluated in the well established double exponential decay model [23, 24]. Here, the fits were performed as global fits, where all signals are taken into account at once and the fit for the bound complexes is fitted as a sum of the single contributions. Results of the fits are summarized in Table 1, the error noted is the error due to the fit routine. For both global fits the coefficient of determination is R^2^ = 0.99. The fit analysis for the decays yields that the two time constants in the exponential decays, *τ*_1_ and *τ*_2_, are only slightly different for all proteins and bound proteincomplexes. The recorded signals consist of a fast and a slow decay with *τ*_1_ *≈* (2.0 *±* 0.1) ns and *τ*_2_ *≈* (8.0 *±* 1.0) ns for all proteins.

**Table 1.** Results of the double exponential fit to the different protein time correlated single photon signals at an excitation wavelength of 260 nm.

| | $\tau_1$ (ns) | $\tau_2$ (ns) |
| --- | --- | --- |
| hACE2 | $2.0 \pm 0.1$ | $8.0 \pm 1.4$ |
| RBD | $1.9 \pm 0.1$ | $9.0 \pm 1.9$ |
| S1 | $2.1 \pm 0.1$ | $6.9 \pm 1.5$ |

**Fig 5.**
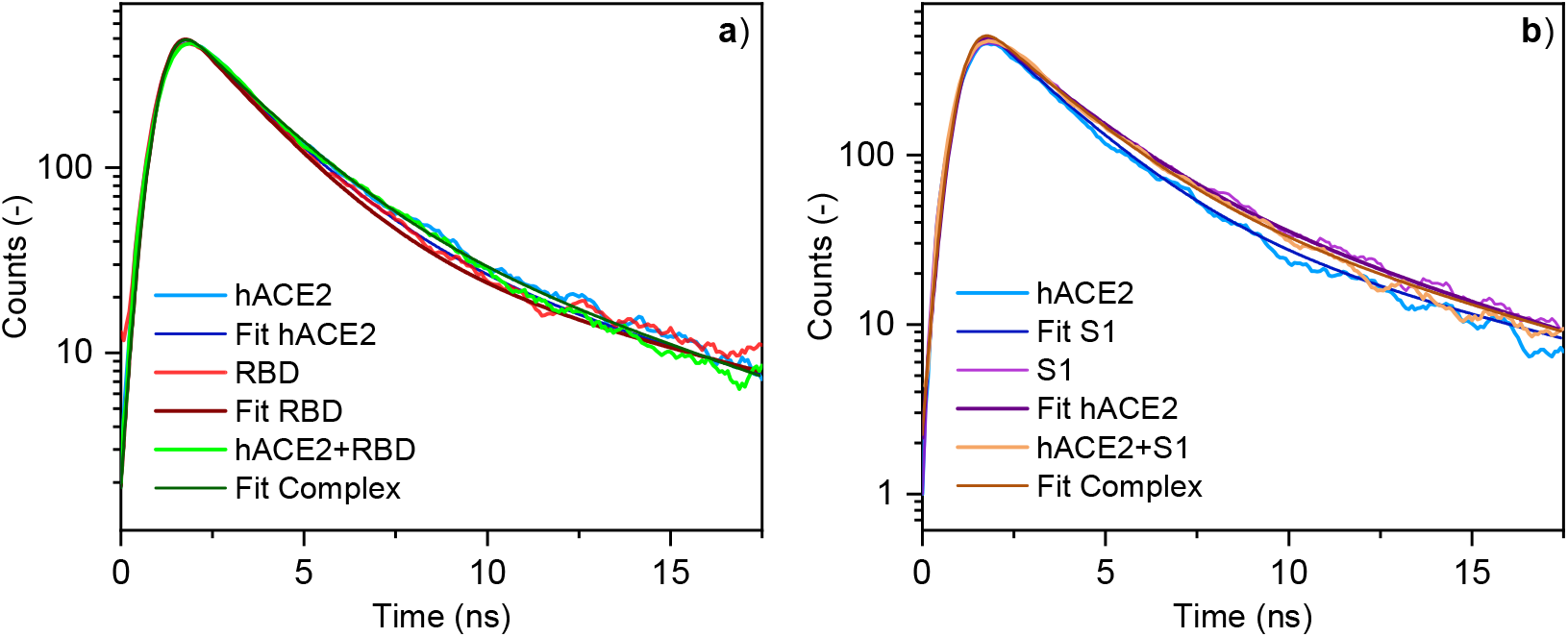
Time correlated single photon counting signals for the different solutions and their respective double exponential decay fit. a) for the RBD case, b) for the S1 case.

## 4 Summary

We have presented fluorescence data for the S1 spike protein of the SARS-CoV-2, its receptor-binding domain and for its most important receptor in human organisms, hACE2. The spectra show distinguishable features such as slight, but clear shifts in the shapes of the LIF signatures for all proteins, e.g. manifested in different FWHMs of the respective fluorescence peaks. Additionally, the signals vary with the excitation wavelength and show for example a clear blueshift of about 5 nm for the receptor-binding domain at *λ*_*ex*_ = 220 nm in contrast to *λ*_*ex*_ = 280 nm. The fluorescence decay times have been determined to be *τ*_1_ *≈* (2.0 *±* 0.1)ns and *τ*_2_ *≈* (8.0 *±* 1.0)ns. For further insight, the limitations are the signal to noise ratios and the maximum possible integration times in order to avoid photolytic decomposition of the proteins in the UV. The data are publicly available for external R&D as described in section 4.

## Data and Materials Availability

Measurement data will be available on biorxiv.org.

## List of Figures

1. Excitation-emission fluorescence map for hACE2 (left), the receptor-binding domain (center) and SARS-CoV-2 S1 (right). Data are available on biorxiv.org as supplementary material.
2. Detailed, normalized fluorescence signals of the three proteins at *λ*_*ex*_ = 220*nm* (left) and *λ*_*ex*_ = 280*nm* (right).
3. Fluorescence spectra of bound protein complexes and their single contributions: a) Receptor-binding domain of the S1 protein and b) the full S1 protein.
4. Differences between the added single contributions and the bound complex for the RBD and the S1 case.
5. Time correlated single photon counting signals for the different solutions and their respective double exponential decay fit. a) for the RBD case, b) for the S1 case.

## List of Tables

1. Results of the double exponential fit to the different protein time correlated single photon signals at an excitation wavelength of 260 nm.

## References

[1] P. Zhou, X.-L. Yang, X.-G. Wang, et al., “A pneumonia outbreak associated with a new coronavirus of probable bat origin,” Nature 579, 270–273 (2020).

[2] D. Cucinotta and M. Vanelli, “WHO Declares COVID-19 a Pandemic,” Acta Bio Medica Atenei Parmensis 91, 157–160 (2020).

[3] M. Haghani and M. C. J. Bliemer, “Covid-19 pandemic and the unprecedented mobilisation of scholarly efforts prompted by a health crisis: Scientometric comparisons across SARS, MERS and 2019-nCov literature,” Scientometrics 125, 2695–2726 (2020).

[4] E. de Wit, N. van Doremalen, D. Falzarano, et al., “SARS and MERS: recent insights into emerging coronaviruses,” Nature Reviews Microbiology 14, 523–534 (2016).

[5] G. Guglielmi, “Rapid coronavirus tests: a guide for the perplexed,” Nature 590, 202–205 (2021).

[6] A. Mohammadi, E. Esmaeilzadeh, Y. Li, et al., “SARS-CoV-2 detection in different respiratory sites: A systematic review and meta-analysis,” EBioMedicine 59, 102903 (2020).

[7] A. D. Teklemariam, M. Samaddar, M. G. Alharbi, et al., “Biosensor and molecular-based methods for the detection of human coronaviruses: A review,” Molecular and Cellular Probes 54, 101662 (2020).

[8] B. Alhalaili, I. N. Popescu, O. Kamoun, et al., “Nanobiosensors for the detection of novel coronavirus 2019-nCoV and other pandemic/epidemic respiratory viruses: A review, sars,” Sensors 20, 6591 (2020).

[9] T. Toptan, L. Eckermann, A. E. Pfeiffer, et al., “Evaluation of a SARS-CoV-2 rapid antigen test: Potential to help reduce community spread?,” Journal of Clinical Virology 135, 104713 (2021).

[10] J. Lukose, S. Chidangil, and S. D. George, “Optical technologies for the detection of viruses like COVID-19: Progress and prospects,” Biosensors and Bioelectronics 178, 113004 (2021).

[11] J. R. Lakowicz, Ed., Principles of Fluorescence Spectroscopy, Springer US (2006).

[12] L. R. Dartnell, T. A. Roberts, G. Moore, et al., “Fluorescence characterization of clinically-important bacteria,” PLoS ONE 8, e75270 (2013).

[13] M. S. Ammor, “Recent advances in the use of intrinsic fluorescence for bacterial identification and characterization,” J Fluoresc 17, 455–459 (2007).

[14] S. Cabredo, A. Parra, and J. Anzano, “Bacteria spectra obtained by laser induced fluorescence,” Journal of Fluorescence 17, 171–180 (2007).

[15] A. Alimova, A. Katz, R. Podder, et al., “Virus particles monitored by fluorescence spectroscopy: A potential detection assay for macromolecular assembly,” Photochemistry and Photobiology 80, 41–46 (2004).

[16] M. Fang, W. Diao, B. Dong, et al., “Detection of the assembly and disassembly of PCV2b virus-like particles using fluorescence spectroscopy analysis,” Intervirology 58(5), 318–323 (2015).

[17] M. Rüdt, P. Vormittag, N. Hillebrandt, et al., “Process monitoring of virus-like particle reassembly by diafiltration with UV/vis spectroscopy and light scattering,” Biotechnology and Bioengineering 116, 1366–1379 (2019).

[18] S, G. JL, B. M, et al., “Non-contact, real-time laser-induced fluorescence detection and monitoring of microbial contaminants on solid surfaces before, during and after decontamination,” Journal of Biosensors & Bioelectronics 09(02) (2018).

[19] V. Gabbarini, R. Rossi, J.-F. Ciparisse, et al., “Laser-induced fluorescence (LIF) as a smart method for fast environmental virological analyses: validation on picornaviruses,” Scientific Reports 9, 1–7 (2019).

[20] F. Gebert, M. Kraus, L. Fellner, et al., “Novel standoff detection system for the classification of chemical and biological hazardous substances combining temporal and spectral laser-induced fluorescence techniques*,” The European Physical Journal Plus 133, 1–5 (2018).

[21] M. Carestia, R. Pizzoferrato, M. Gelfusa, et al., “Development of a rapid method for the automatic classification of biological agents’ fluorescence spectral signatures,” Optical Engineering 54, 114105 (2015).

[22] Y. M. Bar-On, A. Flamholz, R. Phillips, et al., “SARS-CoV-2 (COVID-19) by the numbers,” eLife 9 (2020).

[23] K. Willis, A. Szabo, J. Drew, et al., “Resolution of heterogeneous fluorescence into component decay-associated excitation spectra. application to subtilisins,” Biophysical Journal 57, 183–189 (1990).

[24] A. G. Szabo and D. M. Rayner, “Fluorescence decay of tryptophan conformers in aqueous solution,” Journal of the American Chemical Society 102, 554–563 (1980).

